# Anti-SARS-CoV-2 activity of *Andrographis paniculata* extract and its major component Andrographolide in human lung epithelial cells and cytotoxicity evaluation in major organ cell representatives

**DOI:** 10.1101/2020.12.08.415836

**Authors:** Khanit Sa-ngiamsuntorn, Ampa Suksatu, Yongyut Pewkliang, Piyanoot Thongsri, Phongthon Kanjanasirirat, Suwimon Manopwisedjaroen, Sitthivut Charoensutthivarakul, Patompon Wongtrakoongate, Supaporn Pitiporn, Phisit Khemawoot, Somchai Chutipongtanate, Suparerk Borwornpinyo, Arunee Thitithanyanont, Suradej Hongeng

**Affiliations:** Department of Biochemistry, Faculty of Pharmacy, Mahidol University, Bangkok 10400, Thailand; Department of Microbiology, Faculty of Science, Mahidol University, Bangkok 10400 Thailand; Section for Translational Medicine, Faculty of Medicine Ramathibodi Hospital, Mahidol University, Bangkok 10400, Thailand; Excellent Center for Drug Discovery (ECDD), Faculty of Science, Mahidol University, Bangkok, 10400, Thailand; School of Bioinnovation and Bio-Based Product Intelligence, Faculty of Science, Mahidol University, Bangkok 10400, Thailand; Center for Neuroscience, Faculty of Science, Mahidol University, Bangkok 10400, Thailand; Department of Biochemistry, Faculty of Science, Mahidol University, Bangkok 10400, Thailand; ChaoPhya Abhaibhubejhr Hospital, Prachin Buri 25000, Thailand; Chakri Naruebodindra Medical Institute, Faculty of Medicine Ramathibodi Hospital, Mahidol University, Samutprakarn 10540 Thailand; Department of Pediatrics, Faculty of Medicine Ramathibodi Hospital, Mahidol University, Bangkok 10400, Thailand; Department of Biotechnology, Faculty of Science, Mahidol University, Bangkok 10400, Thailand

**Keywords:** SARS-CoV-2, COVID-19, anti-COVID-19, antiviral, *Andrographis paniculata*, andrographolide, cytotoxicity, high-content screening

## Abstract

The coronavirus disease 2019 (COVID-19) caused by a novel coronavirus (SARS-CoV-2) has become a major health problem affecting more than fifty million cases with over one million deaths globally. The effective antivirals are still lacking. Here, we optimized a high-content imaging platform and the plaque assay for viral output study using the legitimate model of human lung epithelial cells, Calu-3, to determine anti-SARS-CoV–2 activity of *Andrographis paniculata* extract and its major component andrographolide. SARS-CoV-2 at 25TCID_50_ was able to reach the maximal infectivity of 95% in Calu-3 cells. Post-infection treatment of *A. paniculata* and andrographolide in SARS-CoV–2 infected Calu-3 cells significantly inhibited the production of infectious virions with the IC_50_ of 0.036 μg/mL and 0.034 μM, respectively, as determined by plaque assay. The cytotoxicity profile developed over the cell line representatives of major organs, including liver (HepG2 and imHC), kidney (HK-2), intestine (Caco-2), lung (Calu-3) and brain (SH-SY5Y), showed the CC_50_ of >100 μg/mL for *A. paniculata* extract and 13.2-81.5 μM for andrographolide, respectively, corresponding to the selectivity index over 380. In conclusion, this study provided experimental evidence in favor of *A. paniculata* and andrographolide for further development as a monotherapy or in combination with other effective drugs against SARS-CoV–2 infection.

## INTRODUCTION

The outbreak of Coronavirus Disease 2019 (COVID-19) is an emergent global health crisis that requires urgent solutions. Since severe acute respiratory syndrome coronavirus 2 (SARS-CoV-2) has been emerged in Wuhan, Hubei, China at the end of 2019,^1^ the total confirmed cases are approaching seventy millions with more than one million deaths globally at the end of December 2020. SARS-CoV–2 is a positive sense, single-stranded, enveloped RNA virus belonging to *Coronaviridae* family, and categorized as a new member of *Betacoronavirus* genus together with severe acute respiratory syndrome coronavirus (SARS-CoV) in 2003 and middle east respiratory syndrome coronavirus (MERS-CoV) in 2012.^2–4^ The host range of SARS-CoV-2 is very broad partly due to the relative conservation of the cellular receptor, angiotensin-converting enzyme 2 (ACE2), among mammals. This phenomenon could explain the interspecies transmission of the virus from animals to cause disease in humans.^5^ Even though the majority of infection was asymptomatic, clinical manifestations of COVID-19 varies widely, ranging from low-grade fever to severe pneumonia and eventually death. The outcome of the infection depended largely on host factors, e.g., age, previous health problems, and immunological status.^6–8^ Among critical manifestations, acute respiratory distress syndrome, cytokine storm, and multi-organ failure are the leading causes of death in COVID-19.^9, 10^

Lacking the effective antivirals against SARS-CoV-2 is undoubtedly one of the main reasons of poor clinical outcomes in severe COVID-19 patients. Drug discovery and repurposing strategies are pursuing to identify potential therapeutic agents.^11–13^ A beneficial instance of this effort was repositioning of remdesivir (that originally developed for Ebola virus infection) for COVID-19 treatment.^14, 15^ Unfortunately, subsequent well-conducted clinical trials revealed that remdesivir had marginally clinical efficacy,^15, 16^ while the cost-effectiveness and accessibility are still a huge concern.^17^ Therefore, further efforts should be made to identify new compounds with the potent anti-SAR-CoV-2 activity. Among all promising candidates, natural products are recognized as the major source for new drug discovery over decades.^18^ Ethnobotanical evidence suggests plant-derived natural compounds are worth investigating to identify potent antivirals against coronaviruses,^19^ while computational approaches have been applied for phytochemicals in order to define their target-specific antiviral potential against SARS-CoV-2.^20^

One of prominent medicinal plants with various pharmacological activities is *Andrographis paniculate*, known as “King of bitters”, which belongs to *Acanthaceae* family.^21^ *A*. *paniculate* is currently used in the traditional medicine to treat common cold, diarrhea, fever due to several infectious causes, and as a health tonic.^22^ A major bioactive component of *A. paniculata* is andrographolide,^23^ a diterpene lactone in the isoprenoid family, which is known for its broad-spectrum anti-viral properties.^24^ Andrographolide has recently predicted *in silico* to have a potent anti-SARS-CoV-2 activity through specific targeting the host ACE2 receptor and the viral factors, i.e., RNA-dependent RNA polymerase, main protease, 3-CL protease, PL protease, and spike protein.^25–28^ Recently, Shi TH, et al., applied an enzyme-based assay to demonstrate an inhibitory effect of andrographolide against SARS-CoV-2 main protease (M^pro^).^29^ Furthermore, our group has utilized a phenotypic cell-based immunofluorescent assay (IFA) to reveal the anti-SARS-CoV-2 effect of *A. paniculata* extract and andrographolide in the African green monkey kidney cells (Vero E6).^30^ Notably that the anti-SARS-CoV-2 activity of *A. paniculata* extract and andrographolide has never been elucidated in the infected human lung epithelial cells.

This study aimed to evaluate the anti-SARS-CoV-2 activity of *A. paniculata* extract and its major component andrographolide by using a legitimate model of infected human lung epithelial cell, Calu-3.^31^ Cytotoxic profiles of *A. paniculata* extract and andrographolide over five major human organs, including lung, brain, liver, kidney, and intestine were achieved by a panel of cell line representatives. The results demonstrated that *A. paniculata* extract and andrographolide have a potent anti-SARS-CoV-2 activity with a high safety margin for major organs in cell culture models.

## MATERIALS AND METHODS

### Cell culture

African green monkey (*Cercopithecus aethiops*) kidney epithelial cells (Vero cells) (ATCC^®^ CCL-81), Vero cell derivative (Vero E6 cells) (ATCC^®^ CRL-1586), human liver cancer cell line (HepG2) (ATCC^®^ HB-8065), human colon cancer cell line (Caco-2) (ATCC^®^ HTB-37), human airway epithelial cell line (Calu-3) (ATCC^®^ HTB-55^)^, human neuroblastoma cell line (SH-SY5Y) (ATCC^®^ CRL-2266) and human normal kidney (HK-2) (ATCC^®^ CRL-2190) were obtained from American Type Culture Collection (ATCC, Manassas, VA, USA). An Immortalized Hepatocyte-Like Cell Line (imHC) was established in-house as previously described^56^ and its hepatic phenotypes were characterized previously.^57, 58^ Vero cells were cultured in Minimum Essential Medium (MEM) (Gibco, Detroit, MI, USA). Vero E6 cells and Caco-2 were cultured in Dulbecco’s Modified Eagle’s Medium (DMEM) (Gibco, Detroit, MI, USA). HepG2, imHC, Calu-3, and SH-SY5Y cells were cultured in Dulbecco’s Modified Eagle Medium:Nutrient Mixture F-12 (DMEM/F-12) (Gibco, Detroit, MI, USA). HK-2 cells were cultured in Dulbecco’s Low Glucose Modified Eagles Medium (DMEM low glucose) (HyClone, Logan, UT, USA). The culture media was supplemented with 10% fetal bovine serum (FBS) (Thermo Scientific Fisher, Waltham, MA, USA) and 100 μg/mL penicillin/streptomycin (Invitrogen, Carlsbad, CA, USA) and 1% GlutaMAX™ (Gibco, Detroit, MI, USA). Cells were incubated at 37°C in a humidified incubator with 5% CO_2_.

### Preparation of SARS-CoV–2 virus

SARS-CoV-2 virus was isolated from nasopharyngeal swabs of a confirmed COVID-19 patient in Thailand (SARS-CoV–2/01/human/Jan2020/Thailand). The virus was propagated in Vero E6 cells as previous described^30^ and stored at −80°C. Viral titration as TCID_50_ titer/mL was performed in the 96-well plate. In brief, the virus stock was titrated in quadruplicate in 96-well plates on Vero E6 cells in serial dilution to obtain 50% tissue culture infectious dose (TCID_50_) using the Reed Muench method.^59^ All the experiments with live SARS-CoV-2 viruses were performed at a certified biosafety level 3 facility at Department of Microbiology, Faculty of Science, Mahidol University.

### Preparation of *Andrographis paniculata* extract and andrographolide

Plant material in this study was common herbs in Thailand, and it was listed in Thai Herbal Pharmacopoeia 2019 (https://bdn.go.th/th/sDetail/10/34/). The plant was identified, authenticated by Chao Phya Abhaibhubejhr Hospital, Prachin Buri, Thailand, and deposited at the herbarium unit. The powder of *Andrographis paniculata* was weighed and soaked in 95% ethanol in a ratio of 1:4. After 24 hours, liquid fraction was separated using a thin straining cloth then filtered through filter paper by vacuum pump. The extract was obtained was concentrated using rotary evaporator at temperature 45°C and then concentrated in water bath at 70°C until it turned concentrated solution. The crude extract was stored at 4°C and protected from light until use.^60^ The andrographolide concentration in the crude extract was measured by HPLC method in following Thai Herbal Pharmacopoeia 2019 protocol. The andrographolide content in crude extract was 7.9% (w/w).^61^ The analytical standard andrographolide was used as a reference (Sigma, St. Louis, MO, USA).

### *In vitro* antiviral assay

Calu-3 cells were seeded at 1×10^4^ cell per well in a 96-black well plate (Corning, NY, USA) and incubated for 24 h at 37°C in 5% CO_2_ atmosphere. Then, culture media was discarded and washed once with phosphate-buffered saline (PBS). Cells were infected with SARS-CoV-2 at 25TCID_50_ for 2 h at 37°C. After viral adsorption, the cells were washed twice with PBS to remove the excessive inoculum, and the fresh culture medium (DMEM/F12 supplemented with 10% FBS, 100 μg/ml Penicillin/Streptomycin) was added. Each concentration of drugs, crude extract, or active compound was inoculated into the culture medium. Infected cells were then maintained at 37°C in 5% CO_2_ incubator for 48 h. Positive convalescent serum (heat-inactivated at 56°C for 30 min) of a COVID-19 patient and anti-human IgG (Santa Cruz Biotechnology, Dallas, TX, USA) were used for a viral inhibition as positive control and negative control, respectively. The experiment was performed in triplicate.

### High-content imaging for SARS-CoV-2 nucleoprotein detection

The cells in the 96-well plate were fixed and permeabilized with 50% (v/v) acetone in methanol on ice for 20 min. The fixed cells were washed once with phosphate-buffered saline with 0.5% Tween detergent (PBST) and blocked with 2% (w/v) BSA in PBST for 1 h at room temperature. Next, the cells were incubated with 1:500 dilution ratio of primary antibody specific for SARS-CoV nucleoprotein^56^ (Rabbit mAb) (Sino Biological Inc. China), which cross-reacting with the NP protein of SARS-CoV–2 as well, for 1 h at 37°C. After incubation, cells were washed with PBST three times. Then, the Goat anti-Rabbit IgG Alexa Fluor 488 (Thermo Scientific Fisher, Waltham, MA, USA) was used as the secondary antibody at 1:500 dilution. Hoechst dye (Thermo Scientific Fisher, Waltham, MA, USA) was applied for nuclei staining. The fluorescent signal was detected and analyzed by the Operetta high-content imaging system (PerkinElmer, Waltham, MA, USA) as previously described.^30^ Percentage of the infected cells in each well was automatically obtained from 13 fields per well using Harmony software (PerkinElmer, Waltham, MA, USA). Data was normalized to the infected control, and IC_50_ value was calculated by GraphPad Prism 7.

### Plaque assay

The viral output in culture supernatants obtained from SARS-CoV-2-infected Calu-3 cells were determined by plaque assay by using Vero cell monolayer. In brief, Vero cells were seeded into 6-well plate 24-h prior to infection. A serial dilution of the virus-containing supernatants was prepared for inoculation into Vero cell monolayer. The cells were incubated for viral adsorption for 1 hr at 37°C incubator and then overlaid with 3 mL/well of MEM medium supplemented with 5% FBS and 1% agarose. The culture was incubated at 37°C in 5% CO_2_ for three days. Thereafter, plaque phenotypes were visualized by staining with 0.33% Neutral Red solution (Sigma-Aldrich, St. Louis, MO, USA) for 5 hrs. Plaque numbers were counted as plaque-forming units per milliliter (PFUs/mL) and reported as the percentage of plaque reduction in comparison to supernatant obtained from the infected Calu-3 without any treatment.

### Cell cytotoxicity assay

All human cell lines were plated in 96-well plates at 5×10^4^ cells/well and treated with various concentrations of *Andrographis paniculata* extract (0-100 μg/mL) and Andrographolide (0-100 μM) for 48 h. Cell viability was examined by an MTT colorimetric assay. In brief, the medium was replaced with MTT [3-(4,5-dimethylthiazol-2-yl)-2,5-diphenyltetrazolium bromide] (Sigma-Aldrich, St. Louis, MO, USA) final concentration at 0.5 mg/mL and incubated for 4 h at 37°C in a humidified incubator with 5% CO_2_. The rest of MTT solution was removed, and formazan crystals were then dissolved with DMSO (Merck, Schuchardt, Darmstadt, Germany). Absorbance was measured at a wavelength of 570 nm by EnVision Multilabel Reader (PerkinElmer, Waltham, MA, USA). Data was normalized to the solvent control, and then CC_50_ values were calculated using GraphPad Prism 7.

## RESULTS AND DISCUSSION

### Optimization of human lung epithelial cells (Calu-3)-based anti-SARS-CoV-2 assay

Urgent demands of the effective anti-SARS-CoV–2 agents draw many attentions of scientific communities to explore for new antiviral candidates. In this study, we aimed to document the anti-SARS-CoV–2 activity of *A. paniculata* extract and andrographolide (**Figure 1A**) for further drug development against COVID-19. Therefore, a robust anti-SARS-CoV-2 screening platform established on the legitimate model is required to facilitate this process. For serving this proposes, we optimized our antiviral assay (which previously established upon Vero E6 cells)^30^ to use Calu-3 human lung epithelial cells as the legitimate host cell for SARS-CoV-2 infection,^31^ and then validated the assay applicability by hydroxychloroquine and niclosamide (**Figure 1B** and **1C**, respectively), two FDA-approved drugs with anti-SAR-CoV-2 activities *in vitro*.^31–33^

**Figure 1.**
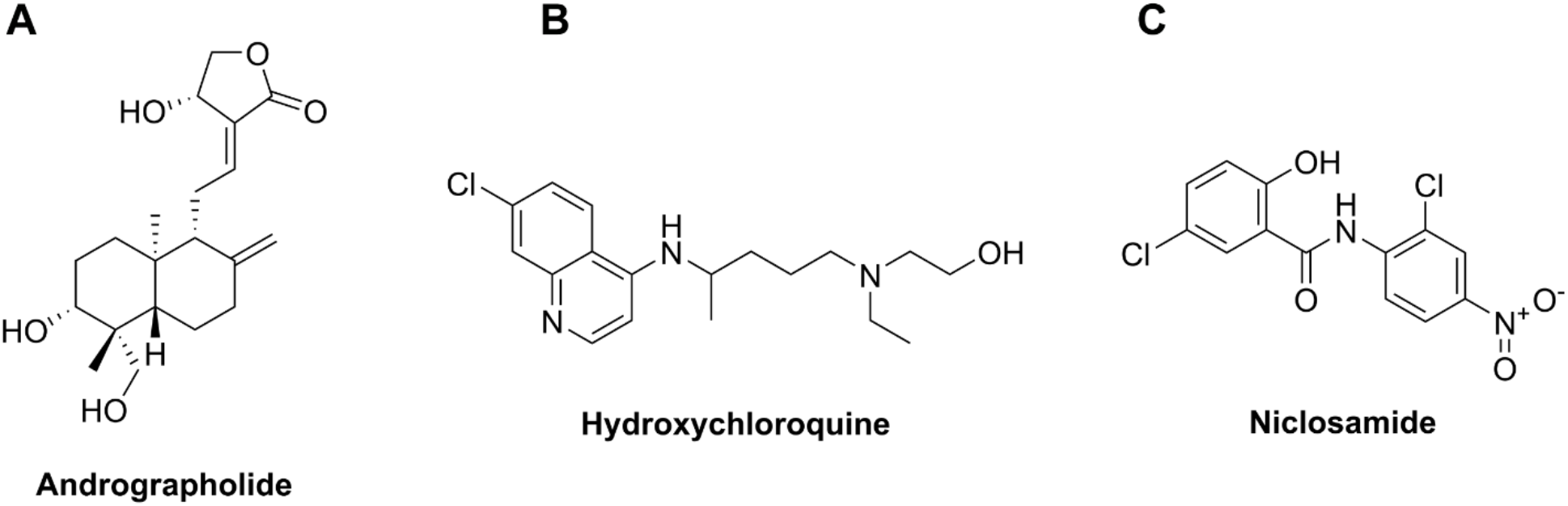
The chemical structures of candidate compounds for screening as anti-SARS-CoV-2 agents. The structure of bioactive compound andrographolide extracted from *Andrographis paniculata* (A). The structures of classical FDA-approved drugs hydroxychloroquine (B) and niclosamide (C) which can exhibit anti-SARS-CoV-2 activity *in vitro*.

Calu-3 cells were cultured until reach confluence, and then infected with various concentrations of SARS-CoV-2, ranging from 0.25TCID_50_, 2.5TCID_50_, and 25TCID_50_ for 2 hrs. The cells were washed twice to remove the excessive inoculum, and further incubated in the fresh culture medium for 48 hrs in the absence or presence of the compounds of interest. To demonstrate the degree of SARS-CoV-2 infectivity, the infected Calu-3 cells were strained by anti-SARS-CoV2 nucleoprotein rabbit monoclonal antibody, following by goat anti-rabbit IgG Alexa Fluor 488, and counted the percentage of fluorescent positive cells by the high-content imaging platform. The percentage of viral infectivity in Calu-3 cells at 48 hours post-infection achieved approximately 1%, 18% and 95%, upon infection at 0.25TCID_50_, 2.5TCID_50_, and 25TCID_50_, respectively (**Figure 2A** and **2B** upper panel). Since SARS-CoV-2 at 25TCID_50_ was able to reach maximal infectivity, it was served as the standard infectious dosage throughout all experiments. To establish the positive control, the neutralizing serum derived from COVID-19 patients with negative SARS-CoV-2 RNA was added to the infected Calu-3 cells. The high-content imaging revealed viral suppression in Calu-3 cells with the percentage of SARS-CoV-2 infectivity of 0%, 10%, 60% and 80% at 1:100, 1:500, 1:2,500 and 1:25,000 dilutions of the neutralizing serum, respectively (**Figure 2A** and **2B** lower panel). In this study, anti-human IgG was served as the negative control.

**Figure 2.**
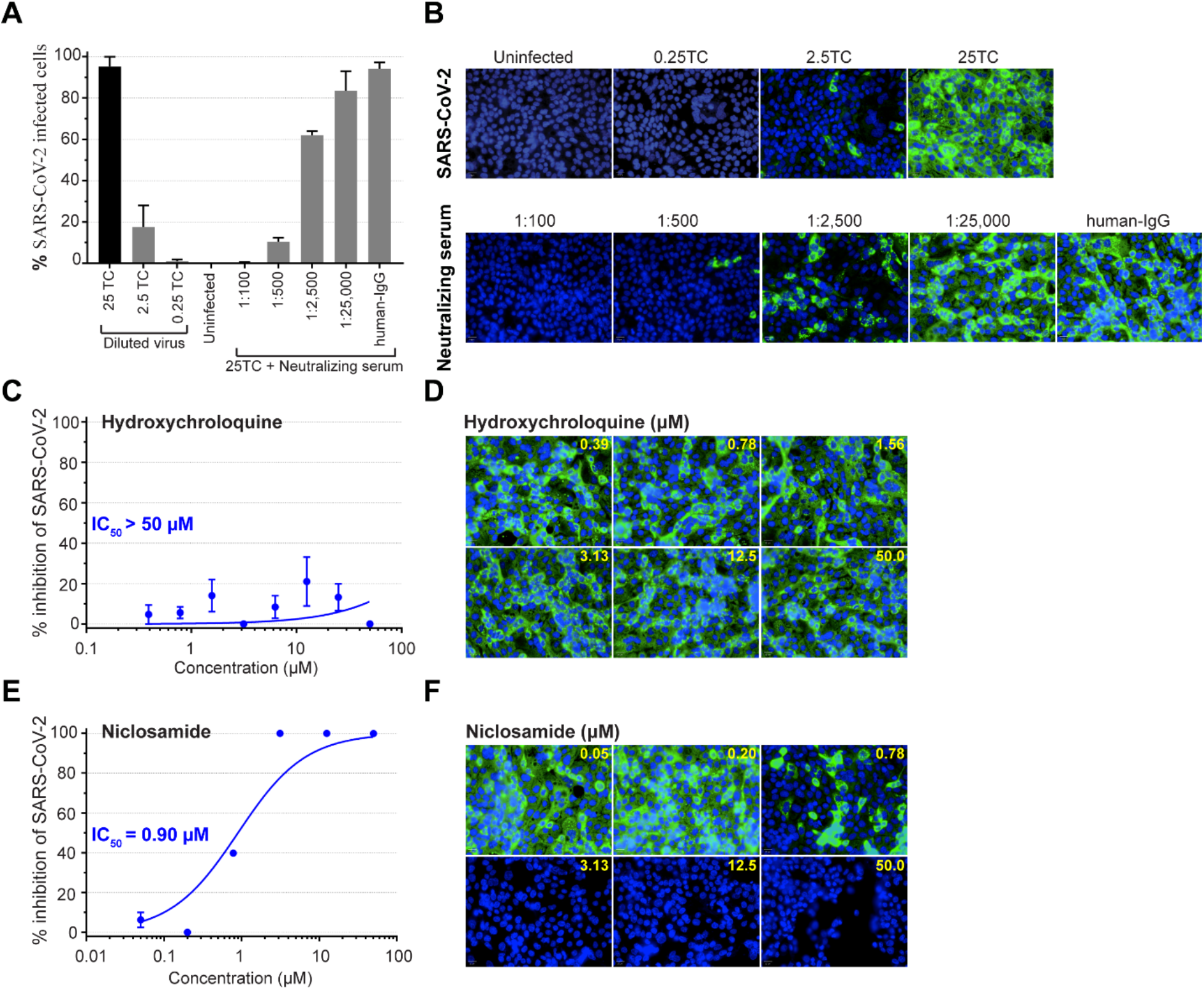
Optimization and validation of anti-SARS-CoV-2 assay using human lung epithelial (Calu-3) cells. (**A**) The optimal dilutions of SAR-CoV-2 TCID_50_, varying from 0.25, 2.5 and 25-fold, were evaluated in Calu-3 cells, in which the 25TCID_50_ showed the maximal infectivity and was used throughout this study. FITC-labeled anti-SARS-CoV nucleoprotein mAb was used to detect the degree of SAR-CoV-2 viral replication. The positive control, neutralizing serum, demonstrated the dose-dependent effect, whereas the negative control human IgG had no antiviral activity. (**B**) Representative fluorescent images show SARS-CoV–2 infectivity at different TCID_50_ (upper row) and anti-SAR-CoV-2 activity of neutralizing serum as compared to human-IgG (lower row). Fluorescent signals: green, anti-SARS-CoV NP mAb; blue, Hoechst. (**C**) Hydroxychloroquine exhibited no effect against SARS-CoV-2 in Calu-3 with IC_50_>50 μM. (**D**) Representative fluorescent images of hydroxychloroquine experiment. (**E**) Niclosamide represented the dose-dependent anti-SARS-CoV-2 activity in Calu-3 with IC_50_ = 0.90 μM. (**F**) Representative fluorescent images of niclosamide experiment.

As aforementioned, hydroxychloroquine and niclosamide^31–33^ were applied to evaluate the validity of Calu-3-based anti-SARS-CoV-2 assay. Hydroxychloroquine, a classical anti-malarial drug, had no inhibitory effect against SARS-CoV-2 infection in human lung epithelial cells (IC_50_>50 μM) (**Figure 2C** and **2D**), while niclosamide, a classical anti-helminthic drug, inhibited SARS-CoV-2 infection with the IC_50_ of 0.90 μM (**Figure 2E** and **2F**). Of note, hydroxychloroquine can exhibit antiviral effect in Vero E6 cells^30^ but not Calu-3 human lung epithelial cells.^31^ These results were consistent to previous reports ^31, 32^ and supported the validity of Calu-3-based anti-SARS-CoV-2 assay used in this study.

### Dose-response relationship of *A. paniculata* extract and andrographolide in SARS-CoV-2 infected human lung epithelial cells

To investigate whether *A. paniculata* extract and its major component andrographolide have potentials for anti-SARS-CoV–2 agents, SARS-CoV–2 infected Calu-3 cells were treated for 48-h with 4-fold dilutions of *A. paniculata* extract (0.05-50 μg/mL) or andrographolide (0.05-50 μM), respectively. The results demonstrated both *A. paniculata* extract (**Figure 3A** and **3B**) and andrographolide (**Figure 3C** and **3D**) inhibited SARS-CoV-2 replication in the dose-dependent manner.

**Figure 3.**
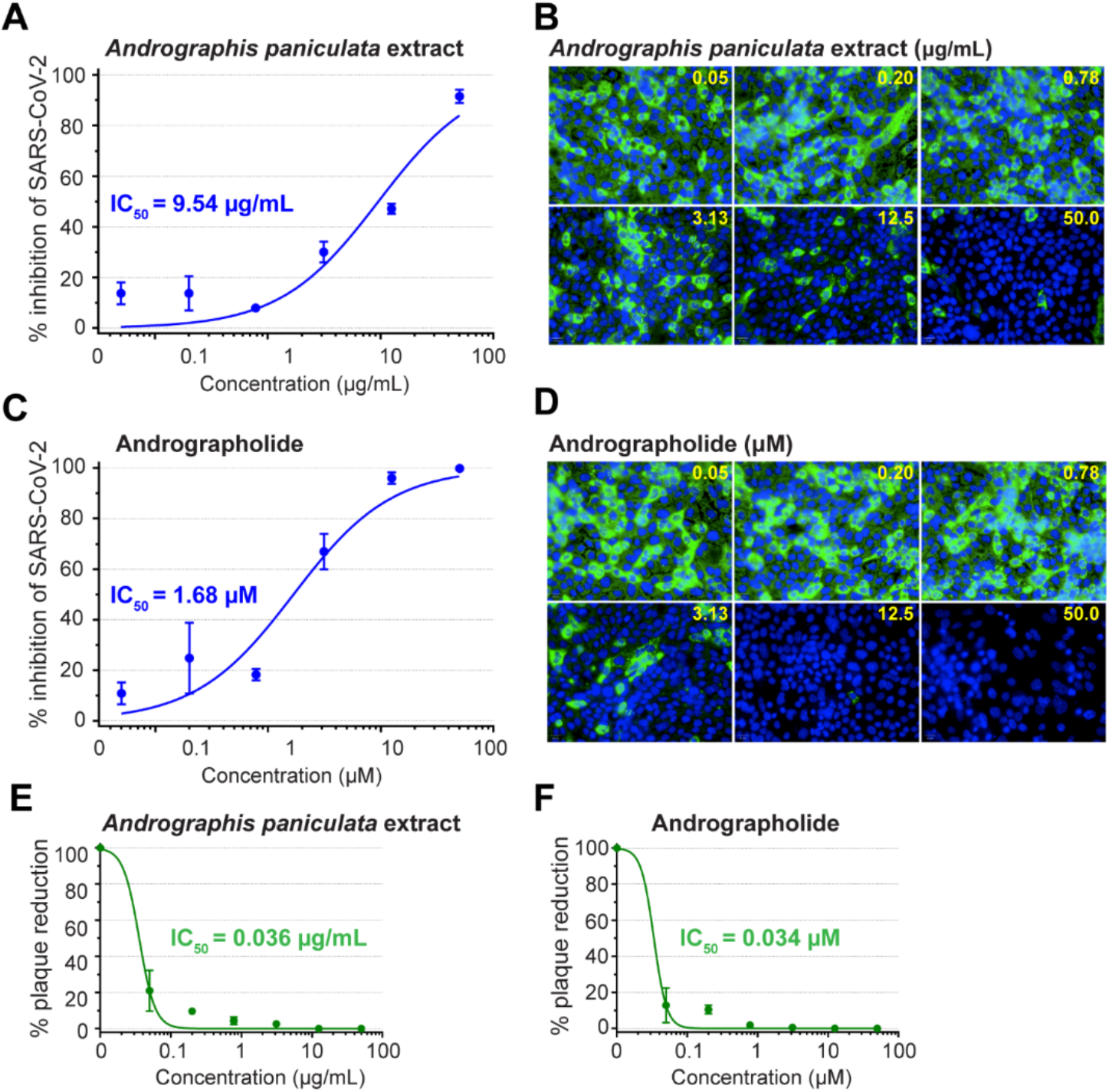
Anti-SARS-CoV-2 activity of *Andrographis paniculata* extract and andrographolide. SARS-CoV-2 infected Calu-3 (at 25TCID_50_) was treated with various concentrations of *A. paniculata* extract or andrographolide for 48 h before harvesting for high-content imaging analysis. (**A**) *A. Paniculata* extract showed the dose-dependent inhibition of SARS-CoV-2 infection. (**B**) Representative fluorescent images of *A. paniculate* experiment. (**C**) Andrographolide, the major component of *A. Paniculata* extract, exhibited potent anti-SARS-CoV-2 activity. (**D**) Representative fluorescent images of andrographolide experiment. Inhibition of infectious virions released from SARS-CoV-2 infected Calu-3 was evaluated by plaque reduction assay after treatment with (**E**) *A. paniculata* extract and (**F**) andrographolide. All experiments were performed in three biological replicates.

To confirm anti-SARS-CoV2 activity of *A. paniculata* extract and andrographolide, analysis of viral output using plaque reduction assay was performed. At 48-h post-infection in the absence or presence of compounds of interest, the culture supernatants were harvested to determine the number of the infectious virion production from SARS-CoV-2 infected Calu-3 cells by plaque assay. From the result, evaluation of viral output was consistent to that of high-content imaging study (**Figure 3A-3D**), in which *A. paniculata* extract and andrographolide again demonstrated the dose-response relationship (**Figure 3E** and **3F**) with the IC_50_ of 0.036 μg/mL for *A. paniculata* and 0.034 μM for andrographolide (**Figure 3E** and **3F**).

It was interesting that the IC_50_ values of *A. paniculata* extract and andrographolide varied between the high-content imaging IFA and viral output study using plaque assay. Taking our previous study^30^ into account, the IC_50_ of *A. paniculata* extract and andrographolide by the types of measure and cell host can be summarized in **Table 1**. The IC50 values of remdesivir^30^ served as the comparator.

**Table 1.**
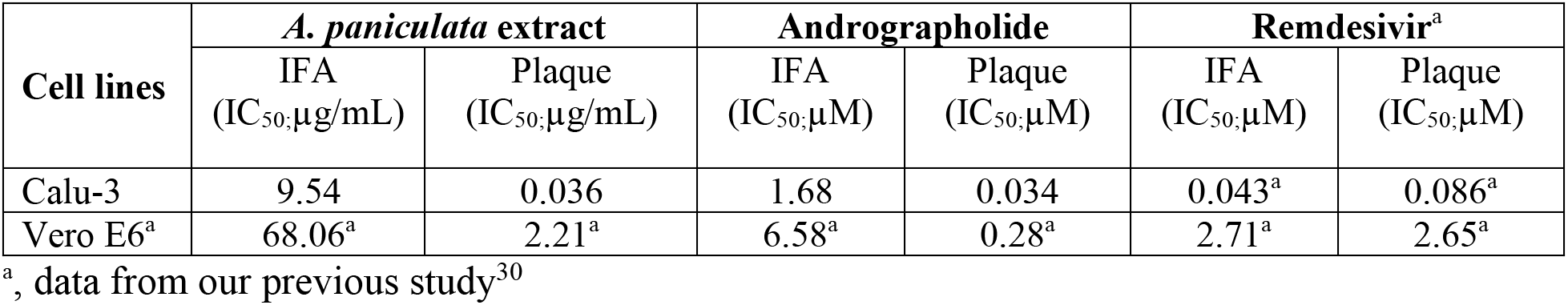
The IC_50_ values of *A. paniculata* extract, andrographolide, and remdesivir evaluated by high-content imaging IFA and plaque assay in Vero E6 and Calu-3.

As shown in **Table 1**, the consistent pattern has been detected for *A. paniculata* extract, andrographolide, and remdesivir; i) the IC_50_ values measured from the assays using Calu-3 were lower than those of Vero E6; ii) the IC50 values measured by plaque assay were lower than those of high-content imaging IFA. The deviation of drug response between two cells of differed species and organs highlighted the importance of using Calu-3 human lung epithelial cells, but not Vero E6 African green monkey kidney cells, as the host cells for *in vitro* SARS-CoV-2 infection experiments.^31^ The dissimilarity of IC_50_ between the IFA and the plaque assay could be explained by their different principle of SARS-CoV-2 detection. The IFA requires the specific antibody to detect the SARS-CoV-2 nucleoprotein derived from the complete virions and sub-viral particles within the host cells, while the plaque assay measures the infectivity of complete virions released from the host cells. Accordingly, it was not surprising that the IC_50_ of the IFA was higher than the plaque assay. Since the IC_50_ of the plaque assay is the standard measure to interpret antiviral potency of the compound of interest, the IC_50_ based on the plaque assay of 0.036 μg/mL for *A. paniculata* and 0.034 μM for andrographolide were applied for data interpretation.

### Cytotoxicity profiles of *A. paniculata* extract and andrographolide

One of the main concerns in medicinal plant-derived drug development is herb-induced injury of the vital organs, especially the liver. To address this issue, six human cell lines which represent five major organs including liver (HepG2 and imHC), kidney (HK-2), intestine (Caco-2), lung (Calu-3) and brain (SH-SY5Y) were applied to evaluate the cytotoxicity profiles of *A. paniculata* extract and andrographolide by MTT assay. The results showed that *A. paniculata* extract had no cytotoxicity to all cell lines examined with the CC_50_ of >100 μg/mL (**Figure 4A-4F**). Considering the antiviral effect with the IC_50_ of 0.034 μM andrographolide (**Figure 3F**), this diterpene lactone showed considerably low-to-no cytotoxic effects on HepG2, imHC, HK-2, Caco-2 and Calu-3 with the CC_50_ of 81.52, 44.55, 34.11, 52.30 and 58.03 μM, and the selectivity index (SI) of 2398, 1310, 1003, 1538 and 1707, respectively (**Figure 4G-4K**). Andrographolide treated SH-SY5Y had the CC_50_ of 13.19 μM and the relatively narrower SI of 388 (**Figure 4L**). The schematic diagram summarized the IC_50_ and the CC50 values of *A. paniculata* extract and andrographolide is shown in **Figure 5**.

**Figure 4.**
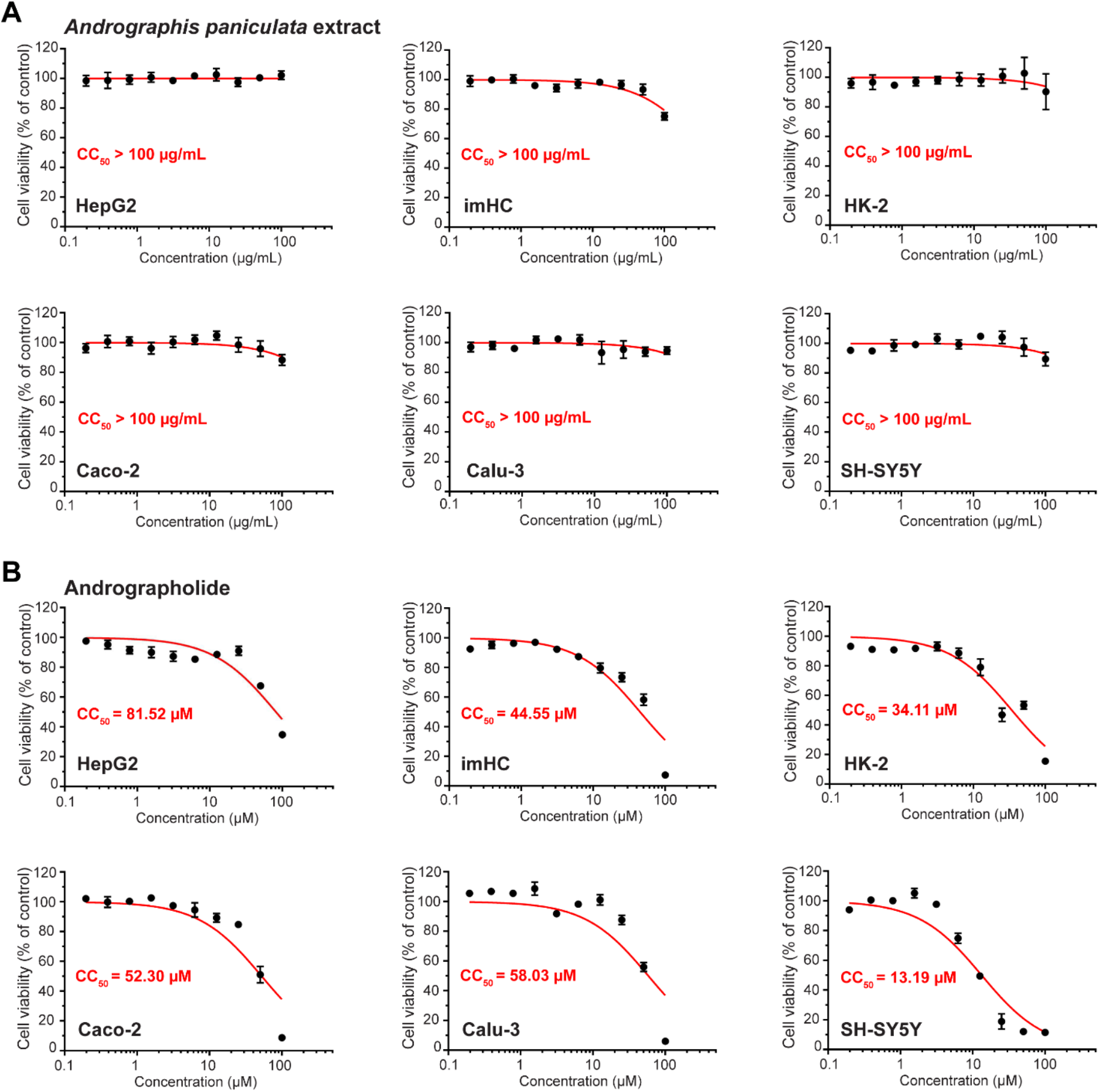
Cytotoxicity profiles of *A. paniculata* extract and Andrographolide over six cell lines representing human major organ. After 48 h treatment of (**A**) *A. paniculata* extract and andrographolide in various cell lines representing liver (HepG2, imHC), kidney (HK-2), intestine (Caco-2), lung (Calu-3), and brain (SH-SY5Y), MTT assay was applied to evaluate the cytotoxicity effect. All experiments were performed in three biological replicates.

**Figure 5.**
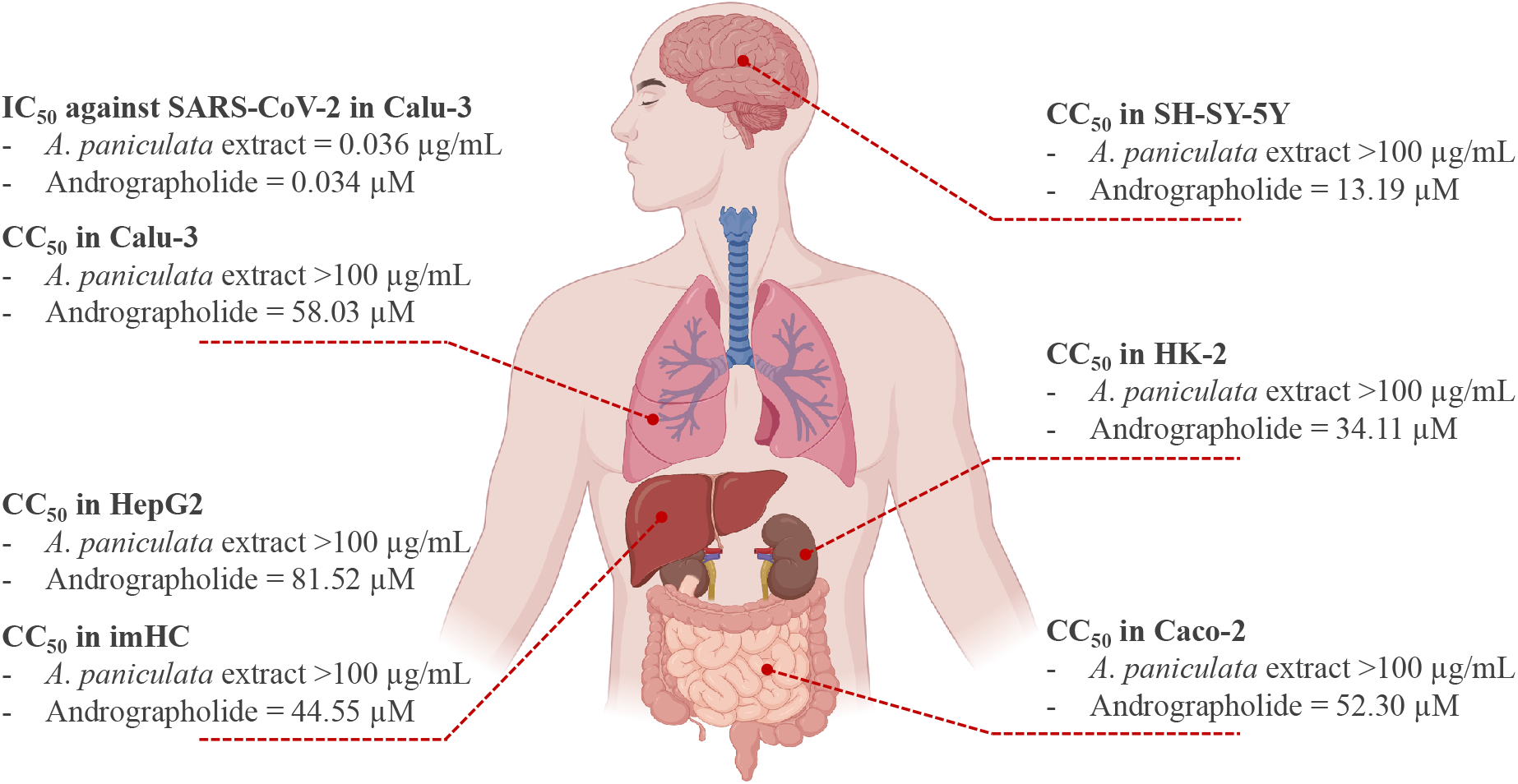
A schematic diagram summarized the IC_50_ and CC_50_ of *A. paniculata* extract and andrographolide across cell line representatives of human major organs.

This finding pointed out that further development of *A. paniculata* extract and andrographolide in preclinical models of COVID-19 should pay attention to the neurologic side effects as well as the amount of andrographolide that pass through the blood brain barrier. In this regard, the BOILED-egg plot using SwissADME program was performed for *in silico* prediction of the probability of compounds being gastrointestinal absorption and barrier penetration.^34, 35^ Based on this prediction, andrographolide is a gastrointestinal absorbable compound without an ability to pass through the blood-brain barrier (**Supplementary Figure 1**). The SwissADME prediction also reveals that andrographolide is a P-glycoprotein (P-gp) substrate,^36^ suggesting that andrographolide would be “pumped out” from central nervous system (CNS) tissues and may not pose a significant neurotoxicity if used in further studies.

### Potential clinical applications of *A. paniculata* extract and andrographolide

*Andrographis paniculata* has classified as an essential plant for traditional medicine in various Asian countries for centuries.^37^ In Thailand, the Ministry of Public Health has registered this plant, so-called Fah Talai Jone, to The National List of Essential Drugs A.D. 1999 (List of Herbal Medicinal Products).^38^ *A. paniculata* extract has long been available in Thailand’s markets as the herbal nutraceuticals in several recipes and brands, in which general population can access and use to treat diarrhea, fever, common cold and viral infection. It is postulated that beneficial effects of *A. paniculata* extract largely depended on its major component andrographolide, the bicyclic diterpene lactone with multi-functionalities including anticancer, antioxidant, anti-inflammatory, immunomodulatory, cardiovascular protection and hepatoprotection, antimicrobial, antiprotozoal.^37, 39–42^

Andrographolide is also well-known for its broad spectrum antiviral properties.^24^ Studies showed andrographolide has been effective against influenza A,^43^ hepatitis C virus,^44^ Chikungunya virus,^40^ HIV,^45^ hepatitis B virus,^46^ Herpes simplex virus 1,^47^ Epstein-Barr virus,^48^ and human papillomavirus.^49^ This study showed that both *A. paniculata* extract and its active component andrographolide had potent inhibitory effect against SARS-CoV-2. This finding opens a possibility to develop *A. paniculata* extract and andrographolide in the context of COVID-19 treatment. In comparison to our previous study,^30^ *A. paniculata* extract and andrographolide exhibited the equivalent IC_50_ against SARS-CoV-2 infection to remdesivir^30^ (**Table 1**). This is an additional rationale to support *A. paniculata* extract and especially andrographolide for further antiviral development.

It has been proposed that andrographolide involves at multiple steps of viral life cycle including viral entry, genetic material replication, protein synthesis and inhibit the expression or function of the mature proteins.^24^ Previous studies suggested andrographolide targeted non-structural proteins of SARS-CoV-2 as the mechanism of action. Enzyme-based assay and *in silico* modelling prediction showed andrographolide could inhibit the main protease (M^pro^) activities of SARS-CoV-2 with the IC_50_ of 15 μM.^26^, ^29^ In addition, Maurya, VK, et al.,^50^ showed that andrographolide has significant binding affinity towards spike glycoprotein of both SARS-CoV-2 and ACE2 receptor and could be develop as a prophylactic agent for limiting viral entry into the host cells. In our study, andrographolide inhibited SARS-CoV-2 at the viral replication and viral release as evidenced from the high-content imaging IFA and viral output study using plaque assay (**Figure 2**). This finding highlighted andrographolide as a potential monotherapy, even though the combinational regimens should be prioritized to increasing the efficacy and reducing the side effects/toxicities.

This study associated with several limitations. Although our findings support andrographolide as a promising candidate for further anti-SARS-CoV-2 development, the low bioavailability of andrographolide might pose a limitation for clinical applications.^51, 52^ Several strategies have been developed to improve andrographolide solubility and bioavailability, i.e., forming complexes with hydroxypropyl-β-cyclodextrin (HP-β-CD),^53^ solid dispersion using a spray-drying technique,^54^ and loading into the nano-emulsion.^55^ Also, it should be noted that this study was conducted using *in vitro* cellular model. The safety and efficacy of andrographolide should be further investigated in preclinical animal models and clinical studies.

In conclusion, this study demonstrated anti-SARS-CoV-2 activity of *A. paniculata* and andrographolide using Calu-3-based anti-SARS-CoV-2 assay. Potent anti-SAR-CoV-2 activities, together with the favorable cytotoxicity profiles, support further development of *A. paniculata* extract and especially andrographolide as a monotherapy or in combination with other effective drugs against SARS-CoV–2 infection.

## ACKNOWLEDGMENTS

We thank Department of Disease Control, Ministry of Public Health Thailand for providing the clinical specimens for viral isolation. This study was supported by the Ramathibodi Research Cluster Grant (CF63010), Faculty of Medicine Ramathibodi Hospital, and Faculty of Science, Mahidol University, Thailand. KS was financially supported by Office of National Higher Education Science Research and Innovation Policy Council through Program Management Unit for Competitiveness (C10F630093). SoC was financially supported by the Faculty Staff Development Program of Faculty of Medicine Ramathibodi Hospital and the Office of National Higher Education Science Research and Innovation Policy Council of Thailand (NXPO; PMU-B). SH was supported by the Ramathibodi Foundation. SB was supported by the Thailand Center of Excellence for Life Sciences (TCELS) Grant (TC-A15/63). AT was supported by the Chaophaya Abhaibhubejhr Hospital Foundation.

## AUTHOR CONTRIBUTIONS

SB, AT, SH initiate the conception. AS, PKh, SoC, SB, AT, SH developed the design. KS, YP, PT, PKa, SM, SiC performed experiments. All authors analyzed and interpreted the data. KS, AS, YP, PK, SiC, SoC prepared figures and tables. KS and AS wrote the first draft of the manuscript. YP, PT, PKa, SM, SiC, PW, SP, PKh, SoC, SB, AT, SH revised the manuscript. SB and AT finalized the manuscript. SH contributed to the overall research strategy. All authors read and approved the final version of the manuscript.

