## Supplementary figure 1 for "Anti-SARS-CoV-2 activity of *Andrographis paniculata* extract and its major component Andrographolide in human lung epithelial cells and cytotoxicity evaluation in major organ cell representatives"

**SUPPLEMENTARY INFORMATION**

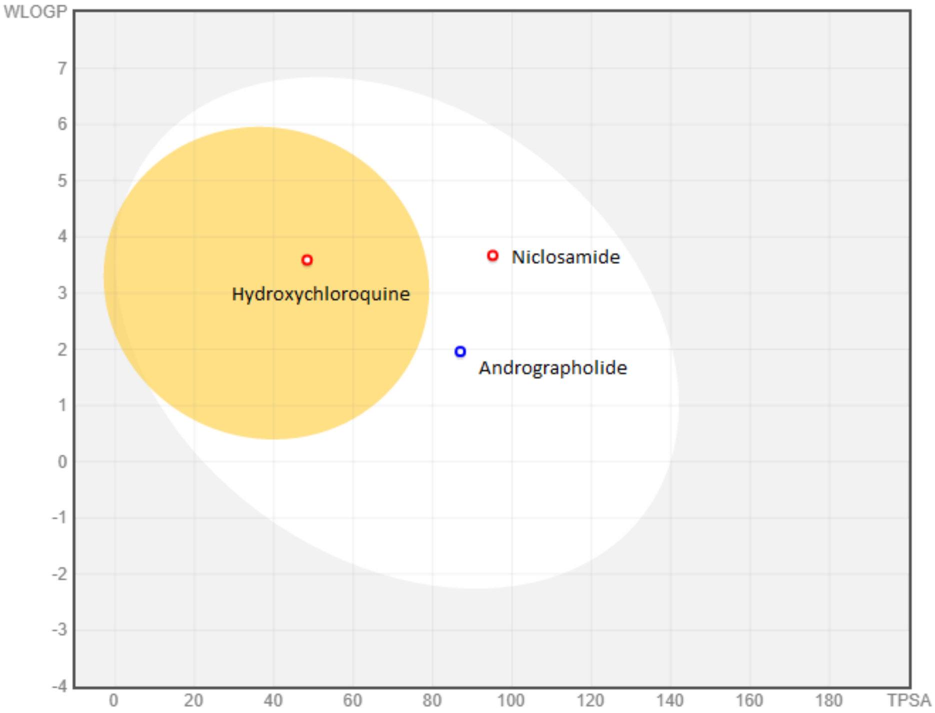


**Supplementary figure 1. The BOILED-egg diagram of andrographolide, niclosamide and hydroxychloroquine predict their gastrointestinal absorption and brain penetration.** Displayed on the X-axis is polarity (tPSA) and Y-axis shows the lipophilicity (WLOGP). The white region is the physicochemical space of molecules with the highest probability of being absorbed by the gastrointestinal tract, and the yellow region (yolk) is for molecules with the highest probability of brain penetration. Yolk and white areas are not completely exclusive. Andrographolide and niclosamide physicochemical characteristics fall in the area outside the space for BBB permeable molecules (white egg). Appeared in the yolk region is hydroxychloroquine. Blue-colored and red-colored circles represent compounds being substrate and non-substrate of the permeability glycoprotein (P-gp), respectively. Of note, P-gp is a member of ATP-binding cassette (ABC) transporters which play an important role in protecting the central nervous system from xenobiotics.[36]
